# Genome sequencing and neurotoxin diversity of a wandering spider *Pardosa pseudoannulata* (pond wolf spider)

**DOI:** 10.1101/747147

**Authors:** Na Yu, Jingjing Li, Meng Liu, Lixin Huang, Haibo Bao, Zhiming Yang, Yixi Zhang, Haoli Gao, Zhaoying Wang, Yuanxue Yang, Thomas Van Leeuwen, Neil S. Millar, Zewen Liu

**Affiliations:** Key laboratory of Integrated Management of Crop Diseases and Pests (Ministry of Education), College of Plant Protection, Nanjing Agricultural University, Weigang 1, Nanjing 210095, China; Novogene Bioinformatics Institute, Beijing 100083, China; Faculty of Bioscience Engineering, Ghent University, Coupure links 653, Ghent B9000, Belgium; Department of Neuroscience, Physiology & Pharmacology, University College London, Gower Street, London WC1E 6BT, United Kingdom

**Keywords:** *Pardosa pseudoannulata*, wandering spider, genome, spidroin, venom, neurotoxin

## Abstract

Spiders constitute an extensive and diverse branch of the phylum Arthropoda. Whereas the genomes of four web-weaver spider species and a single cave-living spider have been determined, similar studies have not been reported previously for a wandering spider. The pond wolf spider, *Pardosa pseudoannulata*, is a wandering hunter that immobilizes prey using venom rather than a web. It is also an important predator against a range of agriculturally important insect pests. The increasing interest in its wandering lifestyle and in the potential of spider venom as a tool for pest control have prompted a detailed study on this wandering spider species. We have generated a high-quality genome sequence of *P. pseudoannulata* and analysed genes associated with the production of silk and venom toxins. Sequencing reveals that *P. pseudoannulata* has a large genome of 4.26 Gb. The presence of only 16 spidroin genes and four types of silk glands is consistent with the moderate use of silk and the lack of a prey-catching web. A large number of genes encode neurotoxins and there is evidence that the majority are highly selective for invertebrates. Comparison between spider species reveals a correlation between spider neurotoxin selectivity for target organisms and spider prosoma size, suggesting a possible coevolution of these two features. The genome data provides valuable insights into the biology of *P. pseudoannulata* and its potential role as a natural enemy in pest control.

## Introduction

Spiders are an important group of arthropods with diverse biological and behavioural characteristics, as is illustrated by their use of both silk and venom to incapacitate prey. Some species of spider build webs for a variety of biological functions, but most notably, for the capture of prey ^1^. In contrast, other species, including the pond wolf spider, *Pardosa pseudoannulata*, have a wandering lifestyle and employ venom for predation and defence (Fig. 1a). The increasing availability of spider genomic and transcriptomic data is helping to provide a better understanding of their biological and evolutionary importance ^2^. In-depth genome sequencing has been reported previously for three Araneoidea spider species that produce prey-catching webs: the velvet spider *Stegodyphus mimosarum* ^3^ and *Stegodyphus dumicola* ^4^, the common house spider *Parasteatoda tepidariorum* ^5,6^ and the golden-orb weaver *Nephila clavipes* ^7^. This has helped to provide information on evolutionary relationships and insights into phenomena such as the diversity of genes that are involved in the production of silk proteins and venom. However, despite the importance and diversity of wandering spiders, only a draft genome has been reported (with 40% coverage) for a sit-and-wait spider (*Acanthoscurria geniculate*, a cave-living species) ^3^.

**Fig. 1.**
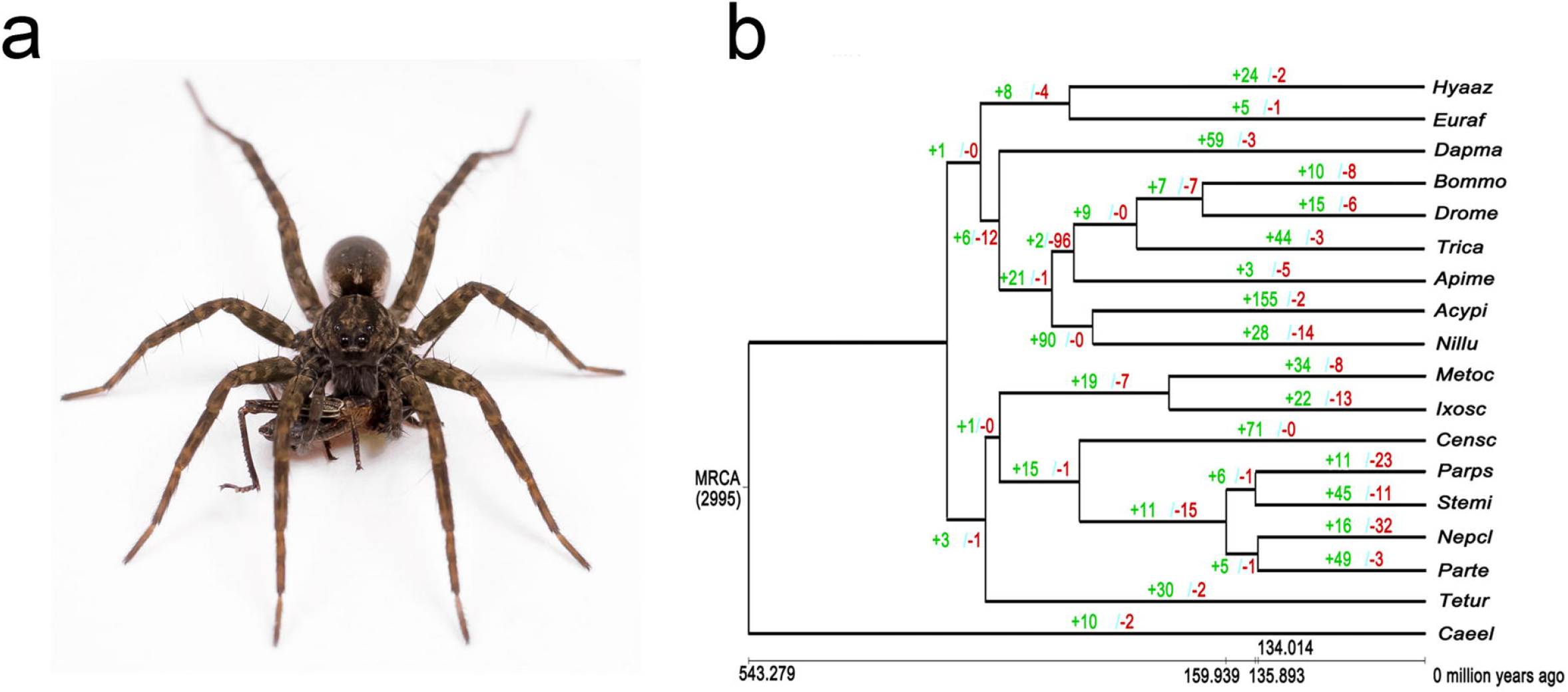
*P. pseudoannulata* and gene gain-and-loss analysis among species of Arthropoda. **(a)** A *P. pseudoannulata* catching prey. **(b)** The divergence time was estimated by PAML mcmctree and is marked with a scale in million years. All internal branches of the tree are 100% bootstrap supported. The numbers next to the branch represent the numbers of expanded (green) and contracted (red) gene families since the split from a most recent common ancestor.

Here, we report the first high-quality genome sequencing of a wandering spider, the pond wolf spider, *Pardosa pseudoannulata*, which belongs to the retrolateral tibial apophysis (RTA) clade of Araneomorphae. An important motivation for undertaking this project was that this would enable a comparison between the genomes of web-building and wandering species, thereby providing insights into their adaptation to differing lifestyles. In addition, *P. pseudoannulata* are predators to a range of insect pests that are of agricultural importance and, as a consequence, *P. pseudoannulata* has been identified as a possible biological control agent. The complete genome sequencing of *P. pseudoannulata* provides a wealth of valuable information, particularly concerning its potential use for insect pest control in integrated pest management.

## Results

High quality genomic DNA was extracted from *P.pseudoannulata* adults for sequencing via Illumina and PacBio technologies. Short-insert (250 bp and 350 bp) paired-end libraries, large-insert (2 kb, 5 kb, 10 kb, 15 kb and 20 kb) mate-pair libraries and 10X Genomics linked-read library were sequenced on the Illumina platform and generated 1771.69 Gb raw data (404.03 × coverage). SMRTbell libraries were sequenced on PacBio Sequel platform and generated 87.37 Gb raw data (19.92 × coverage) (Table S1). Raw data and subreads were filtered. The genome size was estimated via k-mer frequency distribution to be ~4.39 Gb (Table S2). Transcriptome sequencing of four pairs of legs, pedipalp, chelicerae, brain, venom gland, fat body, male silk gland and female silk gland was performed and each generated ~7 Gb raw data (Table S3). A draft genome of 4.27 Gb was eventually assembled with a contig N50 of 22.82 kb and scaffold N50 of 699.15 kb (Table 1, Table S4) with GC content counting for 31.36% (Table S5). The reliability and completeness of the genome assembly were evaluated with EST, CEGMA and BUSCO, with 98.79% of the raw sequence reads aligned to the assembly (Table S6), 84.63% mapped to the genome assembly and 76.71% fully covered by one scaffold with more than 90% of the transcripts mapped to one scaffold (Table S7), 91.53% of conserved eukaryotic genes found (Table S8), and 93.7% of the BUSCO dataset identified in the genome assembly (Table S9).

### Genome statistics and phylogenomics

The assembled genome was ~4.26 Gb with a contig N50 of 22.82 kb and scaffold N50 of 699.15 kb (Table 1). The repeat sequences accounted for ~51.40% of the *P. pseudoannulata* genome and DNA transposons (33.46% of the genome) formed the most abundant category among the TEs, followed by long interspersed elements (LINEs, 3.34%) (Table S10, S11). The gene set contained 23,310 genes, of which 98.8% were supported by homologous evidences or transcriptomic data (Fig S1, Table S12, S13). 19,602 protein-coding genes (accounting for 92.0% of the 21,310 genes) were annotated with at least one public database (Table S14). It was notable that *P. pseudoannulata* genes were generally composed of short exons and long introns, a typical structural feature for Arachnid genes ^3^ (Table S12, S14).

**Table 1.** Summary of the *P. pseudoannulata* genome sequence data.

|  |  |
| --- | --- |
| Total sequencing data | 1,859.06 Gb |
| Sequence coverage | 423.95 x |
| Estimated genome size | 4.42 Gb |
| Assembled genome size | 4.26 Gb |
| Repeat content | 51.39% |
| GC content | 31.36% |
| Number of contigs | 1,242,313 |
| N50 contig size | 22.82 kb |
| Largest contig | 650 kb |
| Number of scaffolds | 1,041,65 |
| N50 scaffold size | 699.15 kb |
| Largest scaffold | 8,106.74 kb |
| Number of protein-coding genes | 21,310 |
| Annotated gene number | 19,602 |
| Average exon length | 179.39 bp |
| Average intron length | 4,132.1 bp |

The phylogenetic relationship of *P. pseudoannulata* with 17 other selected species (Table S15) was analysed using 190 single-copy gene families. According to the phylogenetic analysis, *P. pseudoannulata* diverged from the common ancestral of *S. mimosarum* approximately 135.9 million years ago (MYA) and the lineage of *P. tepidariorum* and *N. clavipes* diverged from the lineage leading to *P. pseudoannulata* and *S. mimosarum* ~159 MYA (Fig. 1b, Fig. S2). The placing of spiders, ticks, scorpions and mites supported the polyphyletic nature of the Acari ^3,8^.

### Gene family expansion

In *P. pseudoannulata* genome, 11 gene families showed significant expansion and 23 gene families showed significant contraction (Fig. 1b). Enrichment of GO terms and KEGG for *P. pseudoannulata* expanded families were performed with the EnrichPipeline ^9^. A false discovery rate (FDR) threshold of 0.05 was used to define GO terms and KEGG that were significantly enriched. Predominantly enriched functional categories for these genes contained several metabolic processes and ion binding that may be related to the environmental adaptation for *P. pseudoannulata* (Table S16, S17).

### Spidroin gene and silk gland

*P. pseudoannulata* typically hunt small insects via wandering in the field rather than building prey-catching webs. Therefore, it is of interest to compare the diversity of spidron genes in a wandering spider compared to that in web waver spiders. In total, sixteen *P. pseudoannulata* putative spidroin genes were obtained and spidroins were classified based on their sequence homology to *N. clavipes* spidroins (Fig. 2a, Table S18). Twelve spidroin genes were designated into five spidroin types as the major ampullate (MaSp, 6), minor ampullate (MiSp, 1), aciniform (AcSp, 3), piriform (PiSp, 1) and tubuliform (TuSp, 1) spidroins based on the sequence alignment of the N-terminal domains with those of *N. clavipes*. The other 4 genes were designated as spidron (Sp) due to lack of clear evidence in sequence similarity (Fig. S3, S4). Up to 28 spidrons were catalogued into 7 spidroin types in *N. clavipes* ^7^ and 19 putative spidroin genes were annotated in *S. mimosarum* genome ^3^. The *P. psueodannulata* spidoin gene repertoire lacked the flagelliform and aggregate spidrons, but this was not surprising given that they mainly function in prey-catching webs ^1,10–14^. The fifteen complete spidroin proteins ranged from 573 (AcSp_2727) to 1977 (MaSp_2831) amino acid residues. Typically, these genes contained only one or two exons but up to 12 exons were also present (e.g. Sp_48488). The spidroins all contained the canonical structure of the repeat region (R) flanked by the relatively conserved N-terminal domain (N) and the C-terminal domain (C) (Fig. 2a, Table S18).

Dissection of adult male and female *P. pseudoannulata* identified only four types of silk glands: the major ampullate gland (Ma), minor ampullate gland (Mi), aciniform gland (Ac) and piriform gland (Pi), which were morphologically similar to the reported black widow spider silk glands (Fig. 2b) ^15,16^. Typically seven types of silk glands are present in orb-weaver spiders ^7^ and the three absent silk gland types in *P. psuedoannulata* were the tubuliform glands, flagelliform glands and aggregate glands. The observed differences in spidroin genes, spidroin types and silk gland types between *P. pseudoannulat*a and web-weaver spiders support the conclusion that silk gland types and silk proteins are specialized in tasks involved in a variety of biological activities ^17,18^. However, whether the spidroin genes were lost in wandering spiders or expanded and differenciated in web-weavers requires further investigation.

**Fig. 2.**
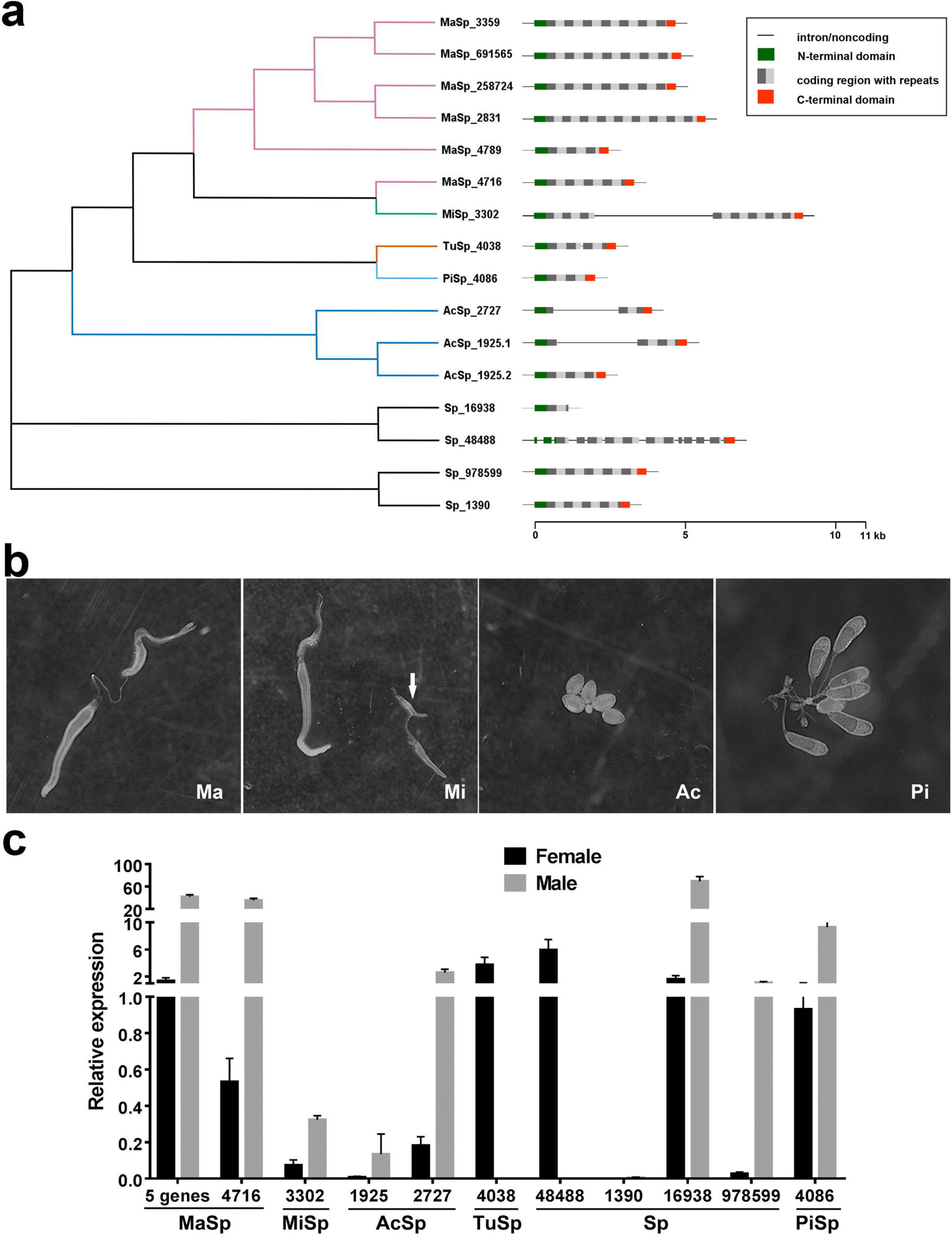
Characterization of spridroin genes in *P. pseudoannulata*. **(a)** Phylogeny and gene structure characteristics of the *P. psedoannulata* spidroins. The phylogenetic tree was constructed with the 130 N-terminal amino acid residues from each putative gene product and transformed cladogram. Gene structures are drawn to scale. The alternated grey and light grey blocks represent only the repeat regions without any sequence preference. Only the N-terminal region and partial repeat region are available for gene MaSp_16938. **(b)** The anatomy of the four types of silk glands in *P. pseudoannulata*. Major ampullate gland (Ma), minor ampullate gland (Mi), aciniform gland (Ac) and piriform gland (Pi) were dissected from a female spider. **(c)** The relative expression of each spidroin gene type in female and male *via* qPCR.

The relative expression of the 16 spidroin genes in female and male *P. pseudoannulata* were quantified. Generally, most spidroin genes were expressed at higher levels in males than in females, however, TuSp_4038 and Sp_48488 were expressed at higher levels in females (Fig. 2c, Table S20). These differences in spidroin expression presumably reflected differences in the requirement for spidroins in each gender. For example, TuSp, which was highly expressed in females, is considered to be the most important component of egg cases ^19–22^. In contrast, PiSp, which was highly expressed in males, may contribute to the silk threads used to attach to substrates ^23,24^. It is also of interest to note that both MaSp and MiSp were abundantly expressed in male *P. psuedoannulata*. Further investigation of the employment of spidroins in male spiders is of interest because, to date, spidroins have been mainly studied in female spiders.

### Venom toxin

Spider venom consists of numerous diverse components, however, in the present study we have focused on the best studied neurotoxins. Spider neurotoxins are peptides that contain 6-14 cysteine residues forming disulphide bridges and typically comprise the inhibitor cysteine knot (ICK) motif ^25^. Thirty-two putative neurotoxin precursor genes have been identified from the venom gland transcriptome and genome analysis, forming six distinct families ^26^. The neurotoxin genes of similar sequence often cluster on the same scaffold (Table S21). Other venom components have also been annotated, including venom allergen 5, hyaluronidase, astacin-like metalloprotease toxins, and Kunitz-type protease inhibitors (Table S22).

To date, at least 260 spider neurotoxins acting on ion channels have been documented in the spider toxin database ArachnoServer 3.0 ^27,28^. With the aim of identifying the possible targets of *P. paseudoannulata* neurotoxins, we conducted a phylogenetic analysis with 29 *P. pseudoannulata* neurotoxins and 48 neurotoxins with known molecular targets from 14 other spider species (Fig. 3a). Further, *P. pseudoannulata* neurotoxins are clustered into groups with neurotoxins that target invertebrates only (15/29), vertebrates only (2/29) and both invertebrates and vertebrates (7/29) according to the documented neurotoxins with known selectivity (designated as reference neurotoxins in Fig. 3a). In addition, a single cluster was identified with no similarity to known neurotoxins (5/29) (Fig. 3a). We expressed one of these neurotoxins U1-lycotoxin-Pp1b *in vitro* and studied its toxicity ^29^. The recombinant U1-lycotoxin-Pp1b did not show significant toxicity to mice at a dose of 10 mg/kg ^30^ but was found to be toxic to the insect *Nilaparvata lugens*, a typical prey of *P. pseudoannulata*, with an *LD*_50_ (medium lethal dose) of 1.874 nmol/g insect (13.70 mg/kg insect) (Fig. S5). Therefore, we classified the five unclustered neurotoxins into the group ‘toxic to invertebrates’, based on the observed selectivity of U1-lycotoxin-Pp1b (Fig. 3b, Table S23). While neurotoxins targeting invertebrates account for 69% (20/29) of genes, their proportion in terms of the transcription level is 93% (Fig. 3c, Table S23). Therefore, *P. pseudoannulata* neurotoxins have high selectivity for insects, consistent with its role as a predator of agricultural insect pests.

Spiders of different body sizes capture prey ranging from small insects to large rodents ^31^. Neurotoxins from *P. psuedoannulata* and 10 other spider species were categorized by their target species as being ‘invertebrate only’, ‘vertebrate only’ or both. Information was retrieved for spider species that have at least 10 neurotoxins with identified targets documented in ArachnoSever 3.0. A clear negative correlation was observed between the percentage of neurotoxins targeting invertebrate only (TX_inv.) and the prosoma length of a spider (liner regression, *F*_1,9_ = 10.17, *P* = 0.011, Percentage of TX_inv.=−2.96 × prosoma length + 89.23, R^2^=0.5304, Fig. 3d, Fig. S6). A correlation between the size of a spider and its prey is likely to be associated with energy and nutritional requirement ^32^. It seems likely that spider neurotoxins have coevolved with spider body size with a shift in prey from smaller invertebrates to larger vertebrates.

**Fig. 3.**
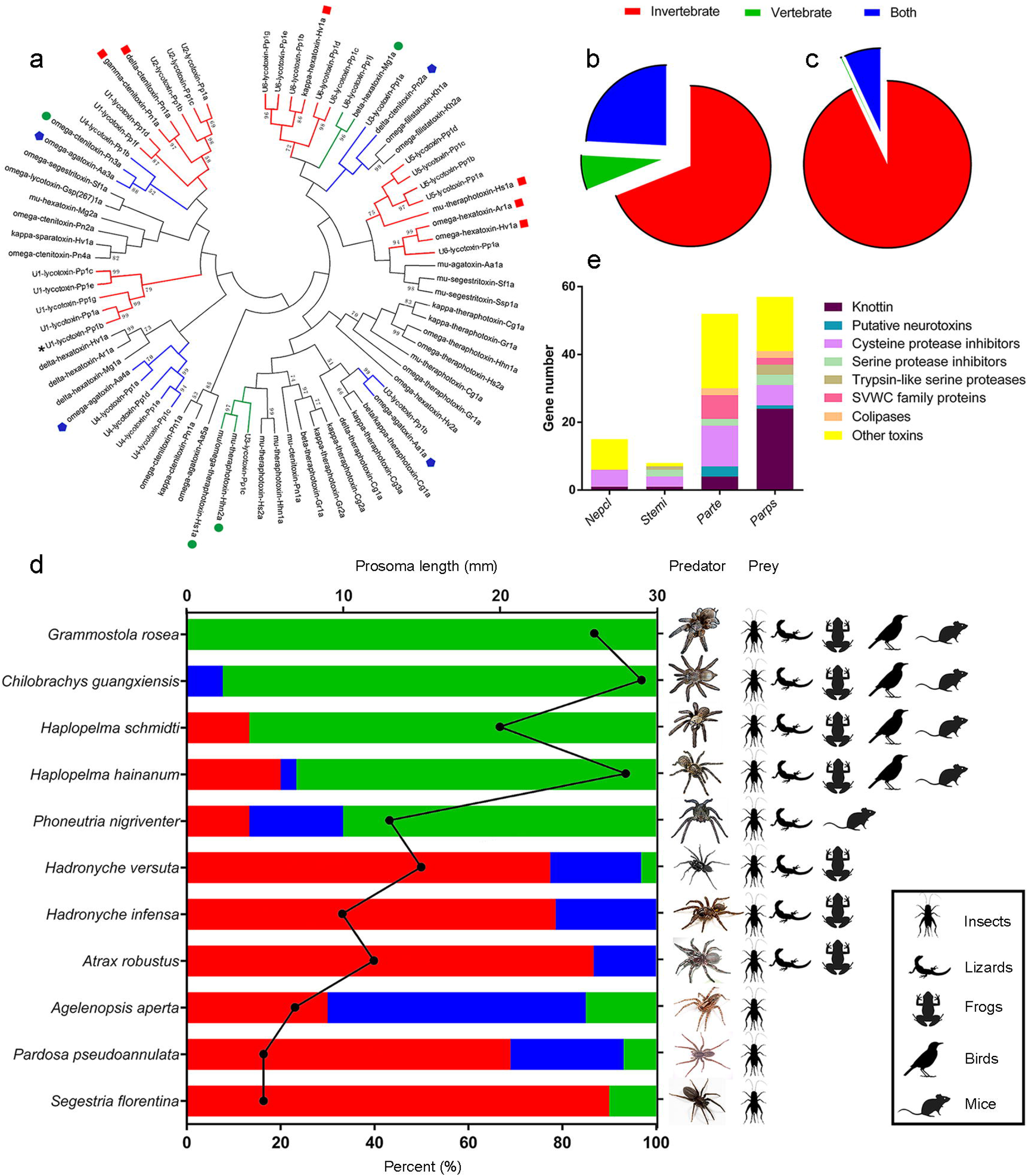
Evolutionary analysis of neurotoxins from *P. pseudoannulata* and other spiders. Neurotoxins are categorized according to their target organism as invertebrate only (red), vertebrate only (green) and both invertebrate and vertebrate (blue). **(a)** Phylogeny of neurotoxins from *P. pseudoannulata* and other spiders. Neurotoxins with known target organisms were used as a reference for target organism prediction and are marked with red squares, green circles and blue pentagons. Branches containing these neurotoxins were shaded accordingly. An asterisk marks U1-lycotoxin-Pp1b. **(b)** Distribution of *P. pseudoannulata* neurotoxin subtypes in terms of gene numbers. **(c)** Expression of different *P. pseudoannulata* neurotoxin subtypes as the FPKM value in the venom gland transcriptome. **(d)** The percentage of invertebrate-selective neurotoxins negatively correlated with spider prosoma length. Horizontal bars represent the percentage of neurotoxins based on their target organism selectivity. The black data points indicate prosoma length of the predator spider. **(e)** Diversity of toxin composition in four spiders.

Toxins showed considerable diversity in the four species of spiders for which genome sequence data are available (Fig. 3e, Table S24). The annotated toxins can be grouped based on their structural domains. Neurotoxic Knottin toxins comprised nearly half of the toxins in *P. pseudoannulata* and their gene number exceeded that in the three web-weavers. The remarkable abundance of Knottin in *P. pseudoannulata* is consistent with the fact that wandering spiders use venom as their main strategy of predation and defence, whereas orb-weaver spiders rely more on low molecular mass compounds and behavioural adaptations (such as prey-catching webs and sticky glue) ^31^.

## Material and Methods

### Genome sequencing

#### Sample preparation and sequencing

For genome sequencing, three batches of *P. pseudoannulata* adults were collected from spiders reared from two individual egg cases (batch #1 of 40 adults from egg case #1; batch #2 of 60 1^st^-instar spiderlings from egg case #2; and batch #3 of 15 5^th^-instar spiderlings from egg case #2). The two egg cases were derived from two females collected at different time points from the same field in Jiangsu (λ 118.638551, φ 32.030345), China.

High quality genomic DNA was extracted using the conventional phenol/chloroform extraction protocol ^33^ and broken into random fragments for whole-genome shotgun sequencing. The genomic DNA was quality-examined with agarose gel electrophoresis and quantified with Qubit™ system. Short-insert (250 bp and 350 bp) paired-end libraries and large-insert (2 kb, 5 kb, 10 kb, 15 kb and 20 kb) mate-pair libraries were prepared using the standard Illumina protocols. All libraries were sequenced on the Illumina HiSeq 2000 platform with paired-end 150 bp and a total of 1,306.48 Gb sequencing data were produced. To promote genome assembly, the technologies of Pacifc Bioscience’s (PacBio’s) single-molecule real-time (SMRT) sequencing and 10x Genomics link-reads were also applied. For PacBio data, SMRTbell libraries were prepared using 20-kb preparation protocols and sequenced on PacBio Sequel platform, which generated 87.37 Gb (19.92x coverage) sequencing data. The 10X Genomics linked-read library was constructed and sequenced on Illumina Hiseq X Ten platform, which generated 465.21 Gb (106.09x coverage) raw reads. For Illunima sequencing, raw data were filtered according to the following criteria: reads containing adapter sequences; reads with >= 10% unidentified nucleotides (N); reads with low-quality bases (Q-value<5) more than 20%; duplicated reads generated by PCR amplification during library construction. For PacBio sequencing, subreads were filtered with the default parameters. All sequence data were summarized in Table S1.

#### Estimation of genome size

Genome size was estimated by analysing the k-mer frequency. The distribution of k-mer values depends on the genome characteristic and follows a Poisson distribution ^34^. A total of 263 Gb high-quality short-insert reads (350 bp) were used to calculate the 17-mer frequency distribution and then estimate the *P. pseudoannulata* genome size using the following formula ^35^: genome size = (total number of 17-mers)/(position of peak depth).

### Transcriptome preparation and sequencing

A batch of adult spiders were collected randomly from a field in Jiangsu (λ 118.638551, φ 32.030345), China. Tissue samples were dissected from these adults as four pairs of legs, pedipalp, chelicerae, brain, venom gland, fat body, male silk gland and female silk gland. Total RNA for each sample was extracted with TRIZOL Reagent (Thermo Fisher Scientific, Waltham, MA, USA). The concentration and purity of the RNA sample was assessed by Nanodrop spectrophotometer (Thermo Fisher Scientific, Waltham, MA, USA) and the integrity was checked by 2100 Bioanalyzer (Agilent, USA). RNA sequencing (RNA-seq) libraries were constructed using the NEBNext® mRNA Library Prep Master Mix Set for Illumina® (New England Biolab, RRID: SCR_013517) according to the manufacturer’s instructions. All libraries were sequenced on the Illumina Hiseq X Ten with paired-end 150 bp. Information for all RNA-seq data was summarized in Table S3.

### Genome assembly

For genome assembly of *P. pseudoannulata*, Platanus (PLATform for Assembling Nucleotide Sequences, RRID: SCR_015531) (version 1.2.4) ^36^ was first used to construct the genome assembly backbone with all Illumina reads. Briefly, Platanus carried out following three steps: (1) all short-insert paired-end reads were used to construct *de Bruijn* graphs with automatically optimized k-mer sizes; (2) all short-insert paired-end reads and large-insert mate-pair reads were aligned to the contigs for scaffolding; (3) paired-end reads were aligned to scaffolds to close the gap. Then GapCloser (RRID: SCR_015026) (version 1.12) ^37^ was used to fill the gaps in intra-scaffold. Subsequently, the PacBio data were used to fill additional gaps with the software PBJelly (RRID: SCR_012091) (version 1.3.1) ^38^ with default parameters. After that, the resulting scaffolds were further connected to super-scaffolds using the 10X Genomics linked-reads by the software fragScaff (version 140324.1) ^39^

The completeness of the genome assembly and the uniformity of the sequencing were evaluated with several approaches. Briefly, BWA (Burrows-Wheeler Aligner, RRID: SCR_010910) ^40^ was used to align high-quality short-insert reads onto the *P. pseudoannulata* genome with parameters of ‘-k 32 -w 10 -B 3 -O 11 -E 4’. Gene region completeness was evaluated with the transcripts assembled by Trinity (version 2.1.1) ^41^. All assembled transcript (length >= 200bp) were align onto the genome by the software BLAT (BLAT, RRID: SCR_011919) ^36^ with default parameters. CEGMA (Core Eukaryotic Genes Mapping Approach, RRID: SCR_015055) ^42^ was used to identify the exon-intron structures. Further, BUSCO (Benchmarking Universal Single Copy Orthologs, RRID: SCR_015008) (v3.0.2) ^43^ was used to assess the genome completeness with a set of 1066 arthropoda single-copy orthologous genes.

### Genome annotation

#### Repetitive element identification

Transposable elements (TEs) were identified with homology alignment and *de novo* prediction. A *de novo* repeat library was built using RepeatModeler (RRID: SCR_015027) (version 1.0.4) ^44,45^, RepeatScout (RRID: SCR_014653) (version 1.0.5) ^46^, and LTR_FINDER (RRID: SCR_015247) (version 1.06) ^47^ with default parameters. Known TEs were idenfied via homology-based prediction using the RepeatMasker (RRID: SCR_012954) (version 4.0.5) ^44^ with default parameters against the RepBase library ^48,49^. In addition, tandem repeats were identified using Tandem Repeats Finder (RRID: SCR_005659) ^50,51^ with parameters “Match=2, Mismatch=7, Delta=7, PM=80, PI=10, Minscore=50, MaxPeriod=2000”.

#### Protein-coding gene prediction

Protein-coding genes were predicted with a combination of homology-based prediction, *de novo* prediction, and transcriptome sequencing-based prediction methods. For the homology-based gene prediction, protein sequences from four species including *Stegodyphus mimosarum, Parasteatoda tepidariorum, Tetranychus urticae* and *Drosophila serrata* ^52^ were aligned to our assembled genome using TBLASTN (RRID: SCR_011822) ^53,54^ with e-value <=1e-5. The BLAST hits were conjoined with the software Solar ^55^. Then GeneWise (RRID: SCR_015054) (version 2.2.0) ^56,57^ was applied to predict gene models based on the alignment sequences. The *de novo* prediction was performed using Augustus (RRID: SCR_008417) (version 3.0.2) ^58,59^, GeneScan (RRID: SCR_012902) (version 1.0) ^60,61^, GeneID (version 1.4) ^62,63^, GlimmerHMM (RRID: SCR_002654) (version 3.0.4) ^64,65^ and SNAP (Semi-HMM-based Nucleioc acid Parser, RRID: SCR_007936) ^66,67^ on the repeat-masked genome. For the transcriptome-based prediction, RNA-Seq data from different tissues including four pairs of legs, pedipalp, chelicerae, brain, venom gland and fat body were aligned to the *P. pseudoannulata* genome using TopHat (RRID: SCR_013035) (version 2.0.13) ^68,69^ and gene strutures were predicted with Cufflinks (RRID: SCR_014597) (version 2.1.1) ^70,71^. In addition, the RNA-Seq data was assembled by Trinity (RRID: SCR_013048) (version 2.1.1) ^41,72^. These assembled sequences were aligned against our assembled genome by PASA (Program to Assemble Spliced Alignment, RRID: SCR_014656) ^73,74^ and generated gene models were used as the training set for the softwares Augustus, GlimmerHMM and SNAP (Semi-HMM-based Nucleic Acid Parser, RRID: SCR_002127) ^67^. Eventually, gene models obtained from all the methods were integrated into a comprehensive and non-redundant gene set with the software EVidenceModeler (EVM, RRID: SCR_014659) ^75,76^.

#### Functional annotation

To obtain functional annotation, all predicted protein-coding sequences in *P. pseudoannulata* genome were aligned to public databases including National Center for Biotechnology Information nonredundant protein (NR) ^77^ and SwissProt ^78,79^. The known motifs and domains were annotated by searching InterPro databases ^80^ including Pfam (RRID: SCR_004726) (version 27.0) ^81,82^, PRINTS (RRID: SCR_003412) (version 42.0) ^83,84^, PROSITE (RRID: SCR_003457) (version 20.89) ^85,86^, ProDom (RRID: SCR_006969) (version 2006.1) ^87,88^, SMART (RRID: SCR_005026) (version 6.2) ^89,90^ and PANTHER (RRID: SCR_004869) (version 7.2) ^91,92^ with the software InterProScan (RRID: SCR_005829) (version 4.7) ^80,93^. Gene Ontology (GO, RRID: SCR_002811) ^94,95^ terms for each gene were obtained from the corresponding InterPro entry. Kyoto Encyclopedia of Genes and Genomes (KEGG, RRID: SCR_012773) databases ^96,97^ were searched to identify the pathways in which the genes might be involved.

### Phylogeny and divergence time estimation

Gene family analysis was performed with 18 species including *Caenorhabditis elegans*, *Drosophila melanogaster*, *Apis mellifera*, *Tribolium castaneum*, *Nilaparvata lugens*, *Acyrthosiphon pisum*, *Bombyx mori*, *Hyalella Azteca*, *Daphnia magna*, *Eurytemora affinis*, *Tetranychus urticae*, *Ixodes scapularis*, *Metaseiulus occidentalis*, *Centruroides sculpturatus, Stegodyphus mimosarum*, *Parasteatoda tepidariorum*, *Nephila clavipes*, and *P. pseudoannulata* (Table S15). Only the longest transcript of a gene was retained as the representative if the gene had alternative splicing isoforms identified. Genes with protein sequences shorter than 30 amino acids were removed. Then, the similarities between genes in all selected genomes were identified using all-versus-all BLASP with an E-value threshold of 1e-7 and all the blast hits were concatenated by the software Solar ^55^. Finally, gene families were constructed using OrthoMCL ^98,99^ with the setting of “-inflation 1.5”. In total, the protein-coding genes were clustered into 29,995 gene families and 190 single-copy orthologs.

The phylogenetic relationship of *P. pseudoannulata* with the other 17 selected species was analysed using the 190 single-copy gene families. Protein sequences of the ortholog genes were aligned using the multiple alignment software MUSCLE with default parameters ^100^. Then the alignments of each family were concatenated into a super alignment matrix and RAxML (version 8.0.19) ^101,102^ was used to reconstruct the phylogenetic tree through maximum likelihood methods with default substitution model-PROTGAMMAAUTO. Divergence times of these species were estimated using the MCMCtree program in PAML ^103,104^ with the parameters of ‘burn-in=10000, sample-number=100,000 and sample-frequency=2’. Calibration points applied in present study were obtained from the TimeTree database (*Drosophila melanogaster*, *Bombyx mori*, *Tribolium castaneum*, *Apis mellifera*, 238~ 377 MYA; *Apis mellifera*, *Tribolium castaneum*, *Bombyx mori*, *Drosophila melanogaster*, *Acyrthosiphon pisum*, *Nilaparvata lugens*, 295~305 MYA; *Caenorhabditis elegans* and other species, 521~581 MYA). ^105,106^.

### Gene family contraction and expansion

Expansion and contraction analysis of orthologous gene families was performed using CAFÉ program (version 2.1) (Computational Analysis of gene Family Evolution, RRID: SCR_005983) ^107,108^. The program uses a random birth and death model to infer changes of gene families along each lineage of phylogenetic tree. Based on a probabilistic graphical model, this method calculates a p-value for transitions between parent and child nodes gene family size over a phylogeny. The gene families were significantly expanded or contracted in the *P. pseudoannulata* genome with a p-value of 0.05.

### Spidroin gene classification and quantification

Multiple rounds of BLAST (RRID: SCR_004870) were run in the genome database to identify putative spidroin genes. The spidroin genes in *N. clavipes* ^7^ and *S. mimosarum* ^3^ were first used as queries for BLAST and the resultant *P. pseudoannulata* spidroin genes were then added into the query repertoire to run further BLAST searches ^7^. The scaffolds containing the putative spidroin genes were then subjected to Augustus (RRID: SCR_008417) ^109^ for gene prediction and the predicted spidroin genes were manually checked for the presence of structural feature, namely, the N-terminal domain, the repeat region and the C-terminal domain. The manually checked spidroin genes were then examined by BLAST in the silk gland transcriptomes of both males and females for their corresponding transcripts. Spidroin genes were confirmed if at least one transcript aligned with 95% identity. The N-terminal domain (130 amino acids) of the spidroins in *P. pseudoannulata* and *N. clavipes* were aligned with ClustalW function and a phylogenetic tree was constructed with maximum-likelihood method (1000 replicates) in MEGA (RRID: SCR_000667, version 7) ^110^ (Fig. S3, S4). Gene structures were drawn to scale in IBS (version 1.0.3) ^111^.

Silk glands were dissected and identified following protocols relating to the western black widow spider ^15,16^. Images were obtained with a portable video microscope (3R-MSA600, Anyty, 3R Eddyteck Corp., China) and contrasted with Photoshop CS6.

Spider adults were collected from the paddy fields in Nanjing (Jiangsu, China), and reared in laboratory conditions for at least two weeks. Spiders were anesthetized with CO_2_ and silk glands were carefully collected. The entire silk glands from 10 females and 15 males were pooled as one sample, respectively. Three samples were prepared for each gender. Each sample was kept in 200 μL RNA*later* (Thermo Fisher Scientific, Waltham, MA, USA) at −80 °C until total RNA extraction.

Total RNA was extracted from silk gland samples with GeneJET RNA Purification Kit (Thermo Fisher Scientific, Waltham, MA, USA) after removing the RNA*later* and eluted with 44 μL nuclease-free water. Genomic DNA was removed with TURBO DNA-*free* Kit (Thermo Fisher Scientific, RRID:SCR_008452) following the manufacturer’s instructions. The quality and quantity of the total RNAs were monitored with NanoDrop spectrophotometer (Thermo Fisher Scientific) and 2% agarose gel electrophoresis. RNA samples were stored at −80 °C. cDNA was synthesized with 2 μg RNA using PrimeScript RT Reagent Kit (TaKaRa, Kyoto, Japan) and then stored at −20 °C. Primers for quantitative real-time PCR (qPCR) were designed using Beacon Designer (version 7.92, PREMIER Biosoft International, CA, USA) (Table S19). Two pairs of universal primers were designed for MaSp (MaSp_691565, MaSp_3359, MaSp_4789, MaSp_258724, MaSp_2831) and AcSp (AcSp_1925.1, AcSp_1925.2), respectively, due to the high similarity of their sequences. Glyceraldehyde-3-phosphatedehydrogenase (GAPDH) and elongation factor 1-alpha (EF1α) genes were selected as the reference genes for spidroin gene quantification. The specificity and efficiency of the primers were validated via standard curves with five serial cDNA dilutions and the melt curve with a temperature range 60-95 °C. qPCR was performed using SYBR Premix Ex Taq Kit (TaKaRa, Kyoto, Japan) following the manufacturer’s instructions on a 7500 Real-Time PCR System (Applied Biosystems, RRID:SCR_005039). Reagents were assembled in a 20 μL reaction containing 10 μL SYBR Premix Ex Taq, 6.8 μL sterile water, 0.4 μL forward primer, 0.4 μL reverse primer, 0.4 μL ROX Reference Dye II and 2 μL cDNA. The reaction program was 95 °C for 30 sec, 40 cycles of 95 °C for 5 sec and 60 °C for 34 sec. No template control (NTC) and no reverse transcriptase control (NRT) were included as negative controls to eliminate the possibilities of reagent contamination and genomic DNA contamination. Each reaction was performed in two technical replicates and three biological samples were tested. Ct values of qPCR were exported from 7500 Real-Time PCR Software (RRID:SCR_014596) (version 2.0.6). The expression levels of target genes were relative to the geometric mean of two reference genes ^112^ following the 2^−ΔCT^ method ^113^. Statistical analyses were performed with GraphPad Prism (RRID: SCR_002798) (version 7).

### Neurotoxin identification and bioassay

#### Identification of neurotoxin genes and comparative analysis among spiders

Neurotoxin candidates were retrieved via BLAST in the genome database with neurotoxins from ArachnoServer 3.0 as queries ^27,28^. They were identified as neurotoxins when the proteins met the criteria such as containing 6-14 cysteine residues and the canonical neurotoxin domains. The neurotoxins were characterized with their signal peptide via SignalP 4.1 Server ^114,115^, propeptide, and the Cys-Cys disulfide bridge pattern. Inhibitor Cystine Knots (ICK) were predicted on the KNOTTIN database^116^. Toxin genes subjected to interspecific comparison were retrieved from the genome annotation with astacin-like metalloprotease toxin excluded. Non-Knottin toxins were subjected to NCBI domain analysis and grouped accordingly, putative neurotoxins with Spider_toxin or toxin_35 domain, cysteine protease inhibitors containing TY (thyroglobulin type I repeats, accession no. cd00191), serine protease inhibitors containing KU (BPTI/Kunitz family of serine protease inhibitors, accession no. cd00109), Trypsin-like serine proteases with Tryp_SPc (Trypsin-like serine protease, accession no. cd00190), SVWC family proteins containing SVWC (single-domain von Willebrand factor type C proteins, accession no. pfam15430), colipases with COLIPASE (Colipases, accession no. smart00023) and the rest designated as other toxins.

#### Construction of neurotoxin expression vector

The open reading frame of neurotoxin U1-lycotoxin-Pp1b was cloned into the prokaryotic expression vector pLicC-MBP-APETx2 ^29,117^. *Kpn* I and *Ava* I were used for double digestion, and primer sequences were: cggggtaccccggaaaatctgtattttcagggcaaggcatgcaccccaaggttttac (forward) and ccctcgagggttaaccgaatagagtcttaatcttgcc (reverse). For the PCR amplification, the high-fidelity PrimerSTAR (TaKaRa, Tokyo, Japan) was used, and the amplification program was 94°C for 3 minutes, followed by 30 cycles of 94°C for 30 seconds, 55°C for 30 seconds, 72°C for 30 seconds, and finally 72°C for 10 minutes. The PCR products were gel-purified using a Gel Extraction Kit (CWBIO, Nanjing, China), ligated into sequencing vector, and sequenced at Genscript Biotechnology Co. Ltd. (Nanjing, China). Plasmid extraction was performed using Mini Plasmid Extraction Kit I (OMEGA, Guangzhou, China), and double digestion using *Kpn* I (TaKaRa, Tokyo, Japan) and *Ava* I (TaKaRa, Tokyo, Japan). After the target gene and vector were recovered, they were ligated with T4 DNA ligase (TaKaRa, Tokyo, Japan) at 4°C overnight. Subsequently, the ligation product was transformed into *Escherichia coli* BL21 strain, and sequenced at Genscript Biotechnology Co. Ltd. (Nanjing, China). A positive clone was selected and the final concentration of 40% glycerol was added for preservation at −80°C.

#### Expression and purification of the recombinant neurotoxin

Positive clones were cultivated in LB liquid medium containing 100 mg/L carbenicillin at 37°C in a constant temperature shaker with 250 r/min for 14h. The culture broth was inoculated in an LB liquid medium containing 100 mg/L carbenicillin at a ratio of 1:100 and cultured with 250 r/min at 37°C for 4h. IPTG was added into the culture at a final concentration of 0.5 mmol/L, 160r/min, and 25°C to induce the expression for 4h. Bacteria were collected after centrifuging at 4000rpm for 10min at 4°C and resuspended with 20mM Tris-HCl (pH 7.4). Then, 100μg/ml lysozyme, 0.1% Trition X-100, 0.5mM PMSF were added to the suspension and the digestion was performed at 37°C for 1 hour. Bacteria were then ultrasonicated and debris were removed by centrifugation with 16,000 rpm for 20min at 4°C, and the supernatant was filtered through a 0.22 μm filter.

The recombinant toxin was purified using AKTA Avant automated protein purification system (GE, Uppsala, Sweden). Initially, the fusion protein was collected by affinity chromatography using nickel column HisTrap HP (GE, Uppsala, Sweden), and then transferred to TEV protease buffer using desalting column HiPrep 26/10 Desalting (GE, Uppsala, Sweden) and digested with TEV protease (Solarbio, Beijing, China) at 16°C for 10h. Next, TEV protease buffer was replaced by PBS using the desalting column. Finally, the protein label was removed by affinity chromatography using nickel column, and the purity of recombinant toxin was detected by SDS-PAGE.

#### Biological activity assay of recombinant neurotoxin

The insecticidal activity of the recombinant neurotoxin against 5^th^-instar *Nilaparvata lugens* nymphs was determined by microinjection ^118^. *N. lugens* nymphs were injected with neurotoxin solutions at a series of concentrations with 3 replicates of 20 individuals per replicate. PBS was injected as control solution. Before injection, the test insects were anesthetized with CO_2_ and each insect was injected with 30nl of recombinant neurotoxin. After injection, the test insects were checked 1h or 12h later. The toxicity of the recombinant neurotoxin against mice was measured by lateral ventricle injection ^119^.

## Discussions

The complete genome sequencing of the wandering spider *P. pseudoannulata* provides valuable insights into the aspects of the biology of wandering spiders, including diversity of spidroin genes and invertebrate-specific neurotoxins which may have potential importance in developing novel pest management strategies.

The evolutionary diversification of spiders has long been discussed. The focus of the debate has been whether the orb web origin is monophyletic or polyphyletic ^1,2,120^. Notably, the cursorial, non-web building spider taxa from RTA clade has proven to be more important than previously thought in the phylogenetic analysis with morphological, behavioural and molecular evidences ^2,120^. In addition to the distinction in silk use, wandering spiders differs from the web weaver spiders in habitat and biotic interaction, which can also promote the diversification ^2^. Therefore, the massive genomic information of *P. pseudoannulata* offers added value to the diversification analysis. Spiders of different ecological niches have evolved their corresponding behaviours and lifestyle. Genome comparison analysis of spiders of different lifestyles or habitats will reveal primary hints for molecular mechanisms of spiders’ evolutionary adaptation.

The present studies were prompted, in part, an interest in the genetic basis for differences in methods of predation by spiders and, in particular, in the use of neurotoxins to incapacitate insect prey. Although further work will be required to understand the molecular targets and mode of actions of *P. psuedoannulata* neurotoxins, it is possible that a better understanding of spider toxins may lead to the development of novel pest control agents applicable to integrated pest management and other potential biomedical applications.

## Supporting information

Figure S1. Venn diagram of gene sets obtained using three prediction methods (de novo, homology-based, RNAseq-based).

Figure S2. Estimation of divergence time.

Figure S3. Phylogenetic tree of 16 P. pseudoannulata spidroins.

Figure S4. Phylogenetic tree of 16 P. pseudoannulata spidroins and 26 complete N. clavipes spidrions.

Figure S5. Neurotoxin U1-lycotoxin-Pp1 nontoxic to mice but toxic to Nillaparvata lugens

Figure S6. Linear regression analysis of spider prosoma length and the number of neurotoxins targeting invertebrate prey species.

## Availability of supporting data and materials

The *P. pseudoannulata* genome has been deposited in GenBank under accession No. SBLA00000000 and transcriptomes have been deposited to NCBI Sequence Read Archive under accession No. SRR8083387-SRR8083398.

## Additional files

**Table S1-S3.** Statistics of genome and transcriptome sequencing

**Table S4-S6.** Genome assembly

**Table S7-S9.** Genome evaluation

**Table S10-S14.** Genome annotation

**Table S15-S17.** Comparative genomes, gene expansion and contraction

**Table S18-S20.** Spidroin gene classification and quantification

**Table S21-S24.** Venom components and neurotoxins

**Figure S1.** Venndiagram of gene sets obtained using three prediction methods (*de novo*, homology-based, RNAseq-based).

**Figure S2.** Estimation of divergence time.

**Figure S3.** Phylogenetic tree of 16 *P. pseudoannulata* spidroins.

**Figure S4.** Phylogenetic tree of 16 *P. pseudoannulata* spidroins and 26 complete *N. clavipes* spidrions.

**Figure S5.** Toxicity assay of the recombinant neurotoxin U1-lycotoxin-Pp1.

**Figure S6.** Linear regression analysis of spider prosoma length and the number of neurotoxins targeting invertebrate prey species.

## Abbreviations

Ac: aciniform
bp: base pair
BUSCO: benchmarking universal single-copy orthologs
BWA: Burrows-Wheeler Alignment tool
CDS: coding sequence
CEGMA: core eukaryotic genes mapping approach
EF1α: elongation factor 1-alpha
EST: expressed sequence tag
FDR: false discovery rate
FPKM: Fragments Per Kilobase of exon model per Million mapped fragments
GAPDH: glyceraldehyde-3-phosphatedehydrogenase
Gb: gigabases
GO: Gene Ontology
KEGG: Kyoto Encyclopedia of Genes and Genomes
Ma: major ampullate
Mi: minor ampullate
MYA: million years ago
NJ: neighbour joining
NRT: no reverse transcriptase control
NTC: no template control
Pi: piriform
RTA: retrolateral tibial apophysis
Sp: spidroin
TE: transposable element
Tu: tubuliform

## Animal care

Animal experimental procedures were approved by the Laboratory Animal Ethical Committee of Nanjing Agricultural University (No. PZ2019021) and performed accordingly.

## Authors’ contributions

Z. Liu initiated and supervised the project. H. Bao and Y. Yang contributed to the sample collection and handling. M. Liu performed genome sequencing, assembly and primary annotation. M. Liu, J. Li and H. Gao performed the comparative genomic analysis. N. Yu, Y. Zhang, H. Bao, T. Van Leeuwen, N. S. Millar and Z. Liu contributed to the data mining and analysis. N. Yu and Z. Yang conducted the analysis and experimental validation of spidroin genes. L. Huang and Z. Wang performed the analysis and experimental validation of neurotoxins. N. Yu and J. Li submitted data to NCBI. N. Yu and Z. Liu wrote the initial draft of the manuscript. N. Yu, T. Van Leeuwen, N. Millar and Z. Liu revised the manuscript. All authors have read and approved the final manuscript.

These authors contributed equally: Na Yu, Jingjing Li and Meng Liu.

## Current address

Yuanxue Yang, Cotton Research Center, Shandong Academy of Agricultural Sciences, Jinan 250100, China

## Acknowledgments

We thank Dr. Huixing Lin (MOE joint international research laboratory of animal health and food safety, Nanjing Agricultural University) for the neurotoxin assay with mice. The work was supported by National Natural Science Foundation of China (grant number 31772185, 31601656, 31701823).

## References

1. Blackledge, T.A. et al. Reconstructing web evolution and spider diversification in the molecular era. Proceedings of the National Academy of Sciences of the United States of America 106, 5229–5234 (2009).

2. Fernandez, R. et al. Phylogenomics, diversification dynamics, and comparative transcriptomics across the spider tree of life. Current Biology 28, 2190–2193 (2018).

3. Sanggaard, K.W. et al. Spider genomes provide insight into composition and evolution of venom and silk. Nature Communications 5 (2014).

4. Liu, S., Aagaard, A., Bechsgaard, J. & Bilde, T. DNA methylation patterns in the social spider, *Stegodyphus dumicola*. Genes 10 (2019).

5. Gendreau, K.L. et al. House spider genome uncovers evolutionary shifts in the diversity and expression of black widow venom proteins associated with extreme toxicity. BMC Genomics 18, 178 (2017).

6. Schwager, E.E. et al. The house spider genome reveals an ancient whole-genome duplication during arachnid evolution. BMC Biology 15 (2017).

7. Babb, P.L. et al. The Nephila clavipes genome highlights the diversity of spider silk genes and their complex expression. Nature Genetics 49, 895–903 (2017).

8. Regier, J.C. et al. Arthropod relationships revealed by phylogenomic analysis of nuclear protein-coding sequences. Nature 463, 1079–1083 (2010).

9. Chen, S. et al. *De novo* analysis of transcriptome dynamics in the migratory locust during the development of phase traits. Plos One 5 (2010).

10. Adrianos, S.L. et al. *Nephila clavipes* flagelliform silk-like GGX motifs contribute to extensibility and spacer motifs contribute to strength in synthetic spider silk fibers. Biomacromolecules 14, 1751–1760 (2013).

11. Hayashi, C.Y. & Lewis, R.V. Molecular architecture and evolution of a modular spider silk protein gene. Science 287, 1477–1479 (2000).

12. Choresh, O., Bayarmagnai, B. & Lewis, R.V. Spider web glue: two proteins expressed from opposite strands of the same DNA sequence. Biomacromolecules 10, 2852–2856 (2009).

13. Jain, D., Amarpuri, G., Fitch, J., Blackledge, T.A. & Dhinojwala, A. Role of hygroscopic low molecular mass compounds in humidity responsive adhesion of spider’s capture silk. Biomacromolecules 19, 3048–3057 (2018).

14. Singla, S., Amarpuri, G., Dhopatkar, N., Blackledge, T.A. & Dhinojwala, A. hygroscopic compounds in spider aggregate glue remove interfacial water to maintain adhesion in humid conditions. Nature Communications 9, 3048–3057 (2018).

15. Jeffery, F. et al. Microdissection of black widow spider silk-producing glands. Journal of Visualized Experiments (2011).

16. Chaw, R.C. & Hayashi, C.Y. Dissection of silk glands in the western black widow Latrodectus hesperus. Journal of Arachnology 46, 159–161 (2018).

17. Vollrath, F. & Knight, D.P. Liquid crystalline spinning of spider silk. Nature 410, 541–548 (2001).

18. Chaw, R.C. et al. Intragenic homogenization and multiple copies of prey-wrapping silk genes in Argiope garden spiders. Bmc Evolutionary Biology 14 (2014).

19. Casem, M.L., Collin, M.A., Ayoub, N.A. & Hayashi, C.Y. Silk gene transcripts in the developing tubuliform glands of the Western black widow, *Latrodectus hesperus*. Journal of Arachnology 38, 99–103 (2010).

20. Garb, J.E. & Hayashi, C.Y. Modular evolution of egg case silk genes across orb-weaving spider superfamilies. Proceedings of the National Academy of Sciences of the United States of America 102, 11379–11384 (2005).

21. Hu, X.Y. et al. Spider egg case core fibers: Trimeric complexes assembled from TuSp1, ECP-1, and ECP-2. Biochemistry 45, 3506–3516 (2006).

22. Jiang, P. et al. Structure, composition and mechanical properties of the silk fibres of the egg case of the Joro spider, *Nephila clavata* (Araneae, Nephilidae). Journal of Biosciences 36, 897–910 (2011).

23. Blasingame, E. et al. Pyriform spidroin 1, a novel member of the silk gene family that anchors dragline silk fibers in attachment discs of the black widow spider, *Latrodectus hesperus*. Journal of Biological Chemistry 284, 29097–29108 (2009).

24. Geurts, P. et al. Synthetic spider silk fibers spun from pyriform spidroin 2, a glue silk protein discovered in orb-weaving spider attachment discs. Biomacromolecules 11, 3495–3503 (2010).

25. Norton, R.S. & Pallaghy, P.K. The cystine knot structure of ion channel toxins and related polypeptides. Toxicon 36, 1573–1583 (1998).

26. Huang, L., Wang, Z., Yu, N., Li, J. & Liu, Z. Toxin diversity revealed by the venom gland transcriptome of *Pardosa pseudoannulata*, a natural enemy of several insect pests. Comparative Biochemistry and Physiology D-Genomics & Proteomics 28, 172–182 (2018).

27. Pineda, S.S. et al. ArachnoServer 3.0: an online resource for automated discovery, analysis and annotation of spider toxins. Bioinformatics 34, 1074–1076 (2018).

28. ArachnoServer spider toxin database. http://www.arachnoserver.org/mainMenu.html.

29. Anangi, R., Rash, L.D., Mobli, M. & King, G.F. Functional expression in Escherichia coli of the disulfide-rich sea anemone peptide APETx2, a potent blocker of acid-sensing ion channel 3. Marine Drugs 10, 1605–1618 (2012).

30. de Lima, M.E. et al. The toxin Tx4(6-1) from the spider Phoneutria nigriventer slows down Na+ current inactivation in insect CNS via binding to receptor site 3. Journal of Insect Physiology 48, 53–61 (2002).

31. Kuhn-Nentwig, L., Stoecklin, R. & Nentwig, W. Venom composition and strategies in spiders: Is everything possible? Advances in Insect Physiology 40, 1–86 (2011).

32. Nentwig, W. & Wissel, C. A comparison of prey lengths among spiders. Oecologia 68, 595–600 (1986).

33. Sambrook, J., Russell, D.W., Sambrook, J. & Russell, D.W. Molecular cloning: A laboratory manual, (Cold Spring Harbor Laboratory Press, 10 Skyline Drive, Plainview, NY, 11803-2500, USA, 2001).

34. Li, R. et al. The sequence and de novo assembly of the giant panda genome. Nature 463, 1106–1106 (2010).

35. Liu, B. et al. Estimation of genomic characteristics by analyzing k-mer frequency in *de novo* genome projects. Quantitative Biology 35, 62–67 (2013).

36. Kajitani, R. et al. Efficient de novo assembly of highly heterozygous genomes from whole-genome shotgun short reads. Genome Research 24, 1384–1395 (2014).

37. English, A.C. et al. Mind the gap: Upgrading genomes with Pacific Biosciences RS long-read sequencing technology. Plos One 7 (2012).

38. English, A.C., Salerno, W.J. & Reid, J.G. PBHoney: identifying genomic variants via long-read discordance and interrupted mapping. Bmc Bioinformatics 15 (2014).

39. Adey, A. et al. In vitro, long-range sequence information for de novo genome assembly via transposase contiguity. Genome Research 24, 2041–2049 (2014).

40. Li, H. & Durbin, R. Fast and accurate short read alignment with Burrows-Wheeler transform. Bioinformatics 25, 1754–1760 (2009).

41. Grabherr, M.G. et al. Full-length transcriptome assembly from RNA-Seq data without a reference genome. Nature Biotechnology 29, 644–652 (2011).

42. Parra, G., Bradnam, K. & Korf, I. CEGMA: a pipeline to accurately annotate core genes in eukaryotic genomes. Bioinformatics 23, 1061–1067 (2007).

43. Simao, F.A., Waterhouse, R.M., Ioannidis, P., Kriventseva, E.V. & Zdobnov, E.M. BUSCO: assessing genome assembly and annotation completeness with single-copy orthologs. Bioinformatics 31, 3210–3212 (2015).

44. Tarailo-Graovac, M. & Chen, N. Using RepeatMasker to identify repetitive elements in genomic sequences. Current protocols in bioinformatics Chapter 4, Unit 4.10-Unit 4.10 (2009).

45. RepeatModeler. http://www.repeatmasker.org/RepeatModeler/.

46. Price, A.L., Jones, N.C. & Pevzner, P.A. De novo identification of repeat families in large genomes. Bioinformatics 21, I351–I358 (2005).

47. Xu, Z. & Wang, H. LTR_FINDER: an efficient tool for the prediction of full-length LTR retrotransposons. Nucleic Acids Research 35, W265–W268 (2007).

48. Bao, W., Kojima, K.K. & Kohany, O. Repbase Update, a database of repetitive elements in eukaryotic genomes. Mobile DNA 6 (2015).

49. RepBase. https://www.girinst.org/server/RepBase/index.php.

50. Benson, G. Tandem repeats finder: a program to analyze DNA sequences. Nucleic Acids Research 27, 573–580 (1999).

51. Tandem Repeats Finder. http://tandem.bu.edu/trf/trf.download.html.

52. Wang, Z. et al. The draft genomes of soft-shell turtle and green sea turtle yield insights into the development and evolution of the turtle-specific body plan. Nature Genetics 45, 701–706 (2013).

53. Altschul, S.F. et al. Gapped BLAST and PSI-BLAST: a new generation of protein database search programs. Nucleic Acids Research 25, 3389–3402 (1997).

54. TBLASTN. https://blast.ncbi.nlm.nih.gov/Blast.cgi.

55. Darriba, D., Taboada, G.L., Doallo, R. & Posada, D. jModelTest 2: more models, new heuristics and parallel computing. Nature Methods 9, 772–772 (2012).

56. Birney, E., Clamp, M. & Durbin, R. GeneWise and genomewise. Genome Research 14, 988–995 (2004).

57. GeneWise. https://www.ebi.ac.uk/~birney/wise2/.

58. Stanke, M., Schoffmann, O., Morgenstern, B. & Waack, S. Gene prediction in eukaryotes with a generalized hidden Markov model that uses hints from external sources. Bmc Bioinformatics 7 (2006).

59. Stanke, M. & Waack, S. Gene prediction with a hidden Markov model and a new intron submodel. Bioinformatics 19, II215–II225 (2003).

60. Salamov, A.A. & Solovyev, V.V. Ab initio gene finding in Drosophila genomic DNA. Genome Research 10, 516–522 (2000).

61. GeneScan. http://genome.dkfz-heidelberg.de/cgi-bin/GENSCAN/genscan.cgi.

62. Parra, G., Blanco, E. & Guigo, R. GeneID in Drosophila. Genome Research 10, 511–515 (2000).

63. GeneID. http://genome.crg.es/software/geneid/index.html.

64. Majoros, W.H., Pertea, M. & Salzberg, S.L. TigrScan and GlimmerHMM: two open source ab initio eukaryotic gene-finders. Bioinformatics 20, 2878–2879 (2004).

65. GlimmerHMM. http://ccb.jhu.edu/software/glimmerhmm/.

66. Korf, I. Gene finding in novel genomes. BMC Bioinformatics 5 (2004).

67. Semi-HMM-based Nucleic Acid Parser. http://korflab.ucdavis.edu/software.html.

68. Trapnell, C., Pachter, L. & Salzberg, S.L. TopHat: discovering splice junctions with RNA-Seq. Bioinformatics 25, 1105–1111 (2009).

69. TopHat. https://ccb.jhu.edu/software/tophat/index.shtml.

70. Trapnell, C. et al. Transcript assembly and quantification by RNA-Seq reveals unannotated transcripts and isoform switching during cell differentiation. Nature Biotechnology 28, 511–515 (2010).

71. Cufflinks. http://cole-trapnell-lab.github.io/cufflinks/.

72. Trinity. https://github.com/trinityrnaseq/trinityrnaseq/wiki.

73. Haas, B.J. et al. Improving the Arabidopsis genome annotation using maximal transcript alignment assemblies. Nucleic Acids Research 31, 5654–5666 (2003).

74. Program to Assemble Spliced Alignments. https://github.com/PASApipeline/PASApipeline/wiki.

75. Haas, B.J. et al. Automated eukaryotic gene structure annotation using EVidenceModeler and the program to assemble spliced alignments. Genome Biology 9 (2008).

76. EVidenceModeler. https://github.com/EVidenceModeler/EVidenceModeler/.

77. National Center for Biotechnology Information. https://www.ncbi.nlm.nih.gov/protein/.

78. Bairoch, A. & Apweiler, R. The SWISS-PROT protein sequence database and its supplement TrEMBL in 2000. Nucleic Acids Research 28, 45–48 (2000).

79. UniProtKB. http://www.uniprot.org/uniprot/.

80. Mulder, N. & Apweiler, R. InterPro and InterProScan: tools for protein sequence classification and comparison. in Comparative Genomics (ed. Bergman, N.H.) 59–70 (Humana Press, Totowa, NJ, 2007).

81. Bateman, A. et al. The Pfam protein families database. Nucleic Acids Research 28, 263–266 (2000).

82. Pfam. http://pfam.xfam.org/.

83. Attwood, T.K., Beck, M.E., Bleasby, A.J. & Parrysmith, D.J. PRINTS - a database of protein motif fingerprints. Nucleic Acids Research 22, 3590–3596 (1994).

84. PRINTS. http://www.bioinf.man.ac.uk/dbbrowser/PRINTS/

85. Hulo, N. et al. The PROSITE database. Nucleic Acids Research 34, D227–D230 (2006).

86. PROSITE Database of protein domains, families and functional sites. http://www.expasy.ch/prosite/.

87. Bru, C. et al. The ProDom database of protein domain families: more emphasis on 3D. Nucleic Acids Research 33, D212–D215 (2005).

88. ProDom. http://prodom.prabi.fr/.

89. Letunic, I., Doerks, T. & Bork, P. SMART 7: recent updates to the protein domain annotation resource. Nucleic Acids Research 40, D302–D305 (2012).

90. Simple Modular Architecture Research Tool. http://smart.embl-heidelber.de/.

91. Mi, H.Y. et al. The PANTHER database of protein families, subfamilies, functions and pathways. Nucleic Acids Research 33, D284–D288 (2005).

92. PANTHER Classification System. http://www.pantherdb.org/.

93. InterPro: protein sequence analysis & classification. http://www.ebi.ac.uk/interpro/.

94. Ashburner, M. et al. Gene Ontology: tool for the unification of biology. Nature Genetics 25, 25–29 (2000).

95. The Gene Ontology Resource. http://geneontology.org/.

96. Ogata, H. et al. KEGG: Kyoto Encyclopedia of Genes and Genomes. Nucleic Acids Research 27, 29–34 (1999).

97. KEGG: Kyoto Encyclopedia of Genes and Genomes. https://www.genome.jp/kegg/.

98. Li, L., Stoeckert, C.J. & Roos, D.S. OrthoMCL: Identification of ortholog groups for eukaryotic genomes. Genome Research 13, 2178–2189 (2003).

99. OrthoMCL. https://orthomcl.org/orthomcl/.

100. Edgar, R.C. MUSCLE: multiple sequence alignment with high accuracy and high throughput. Nucleic Acids Research 32, 1792–1797 (2004).

101. Stamatakis, A. RAxML-VI-HPC: Maximum likelihood-based phylogenetic analyses with thousands of taxa and mixed models. Bioinformatics 22, 2688–2690 (2006).

102. Stamatakis, A., Hoover, P. & Rougemont, J. A rapid bootstrap algorithm for the RAxML web servers. Systematic Biology 57, 758–771 (2008).

103. Yang, Z. PAML 4: Phylogenetic analysis by maximum likelihood. Molecular Biology and Evolution 24, 1586–1591 (2007).

104. Phylogenetic Analysis by Maximum Likelihood. Vol. 2017.

105. Hedges, S.B., Marin, J., Suleski, M., Paymer, M. & Kumar, S. Tree of life reveals clock-like speciation and diversification. Molecular Biology and Evolution 32, 835–845 (2015).

106. TimeTree. http://www.timetree.org.

107. De Bie, T., Cristianini, N., Demuth, J.P. & Hahn, M.W. CAFE: a computational tool for the study of gene family evolution. Bioinformatics 22, 1269–1271 (2006).

108. Computational Analysis of gene Family Evolution. https://sourceforge.net/projects/cafehahnlab/.

109. Augustus web interface. http://bioinf.uni-greifswald.de/augustus/submission.php.

110 Kumar, S., Stecher, G. & Tamura, K. MEGA7: Molecular evolutionary genetics analysis version 7.0 for bigger datasets. Molecular Biology and Evolution 33, 1870–1874 (2016).

111. Liu, W. et al. IBS: an illustrator for the presentation and visualization of biological sequences. Bioinformatics 31, 3359–3361 (2015).

112. Vandesompele, J. et al. Accurate normalization of real-time quantitative RT-PCR data by geometric averaging of multiple internal control genes. Genome biology 3, research0034 (2002).

113. Livak, K.J. & Schmittgen, T.D. Analysis of relative gene expression data using real-time quantitative PCR and the 2(T)(-Delta Delta C) method. Methods 25, 402–408 (2001).

114. Nielsen, H. Predicting Secretory Proteins with SignalP. in Protein Function Prediction: Methods and Protocols, Vol. 1611 (ed. Kihara, D.) 59–73 (2017).

115. SignalP 4.1 Server. www.cbs.dtu.dk/services/SignalP-4.1/.

116. KNOTTIN database. http://www.dsimb.inserm.fr/KNOTTIN/.

117. Addgene. http://www.addgene.org/.

118. Liu, S., Ding, Z., Zhang, C., Yang, B. & Liu, Z. Gene knockdown by intro-thoracic injection of double-stranded RNA in the brown planthopper, Nilaparvata lugens. Insect Biochemistry and Molecular Biology 40, 666–671 (2010).

119. Garcia, E., Rios, C. & Sotelo, J. Ventricular injection of nerve growth-factor increase dopamine content in the striata of MPTP-treated mice. Neurochemical Research 17, 979–982 (1992).

120. Coddington, J.A., Agnarsson, I., Hamilton, C.A. & Bond, J.E. Spiders did not repeatedly gain, but repeatedly lost, foraging webs. Peerj 7 (2019).

