## Supplementary figures and images for "Genome sequencing and neurotoxin diversity of a wandering spider *Pardosa pseudoannulata* (pond wolf spider)"

### Figure S1. Venn diagram of gene sets obtained using three prediction methods (de novo, homology-based, RNAseq-based).

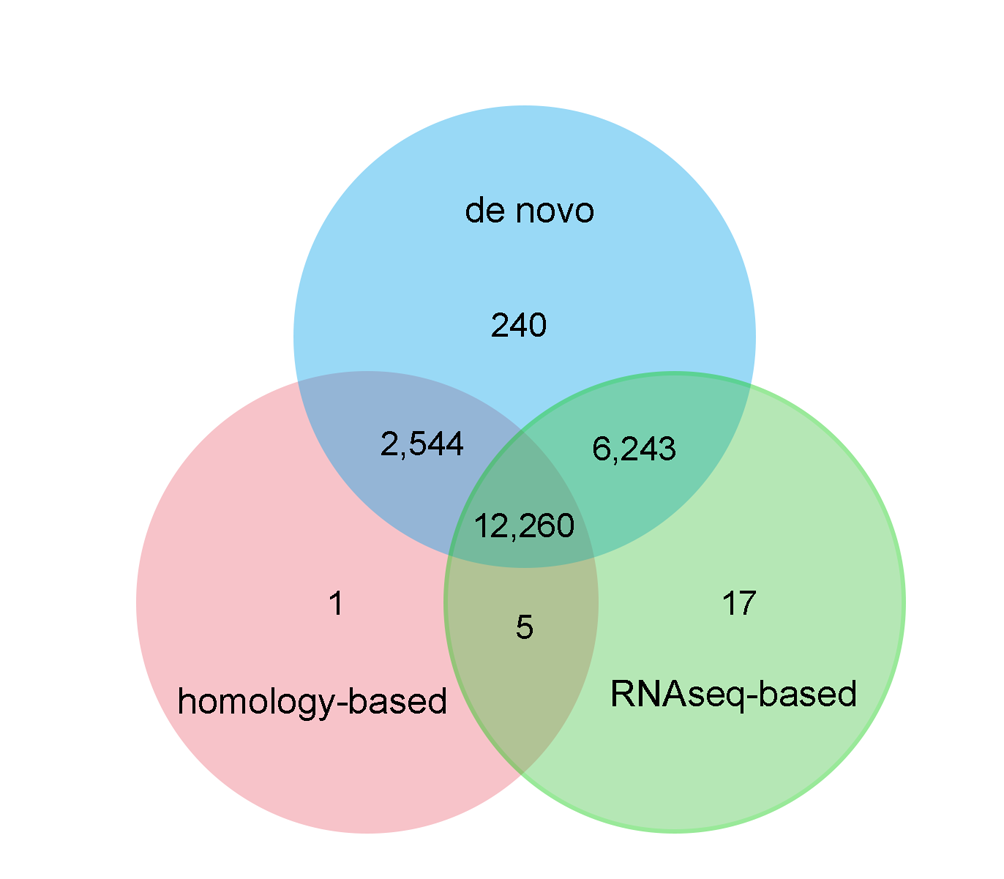

### Figure S2. Estimation of divergence time.

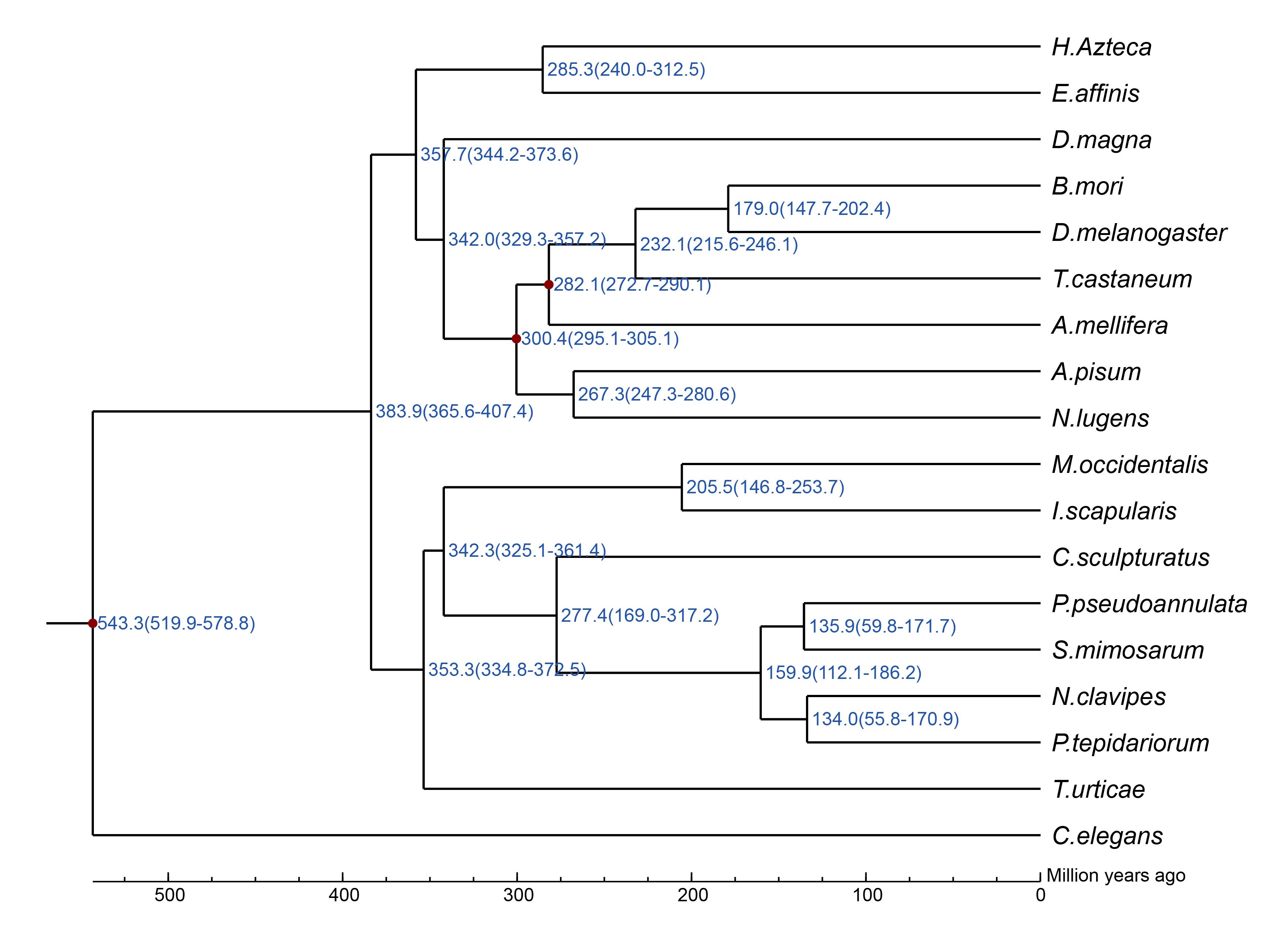

### Figure S3. Phylogenetic tree of 16 P. pseudoannulata spidroins.

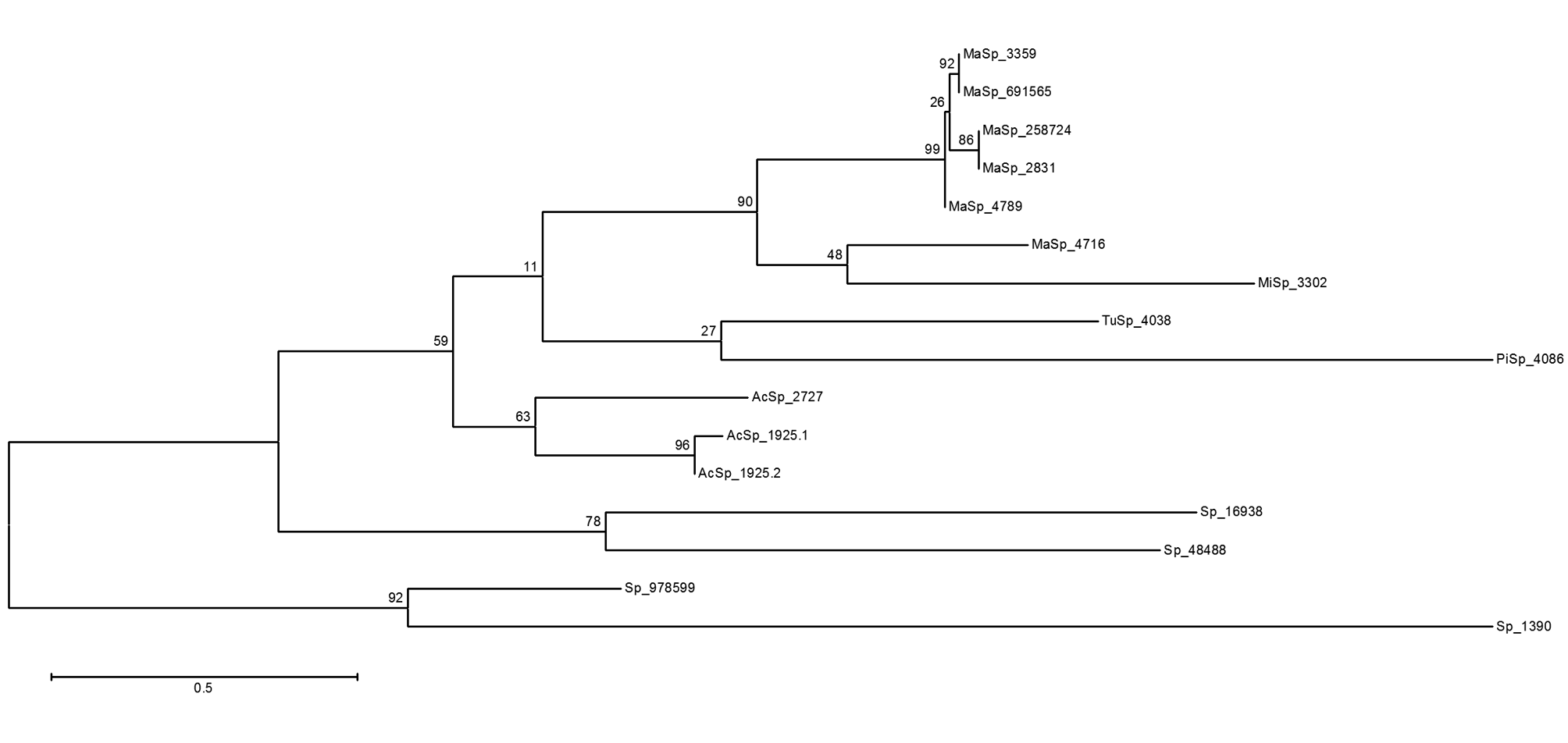

### Figure S4. Phylogenetic tree of 16 P. pseudoannulata spidroins and 26 complete N. clavipes spidrions.

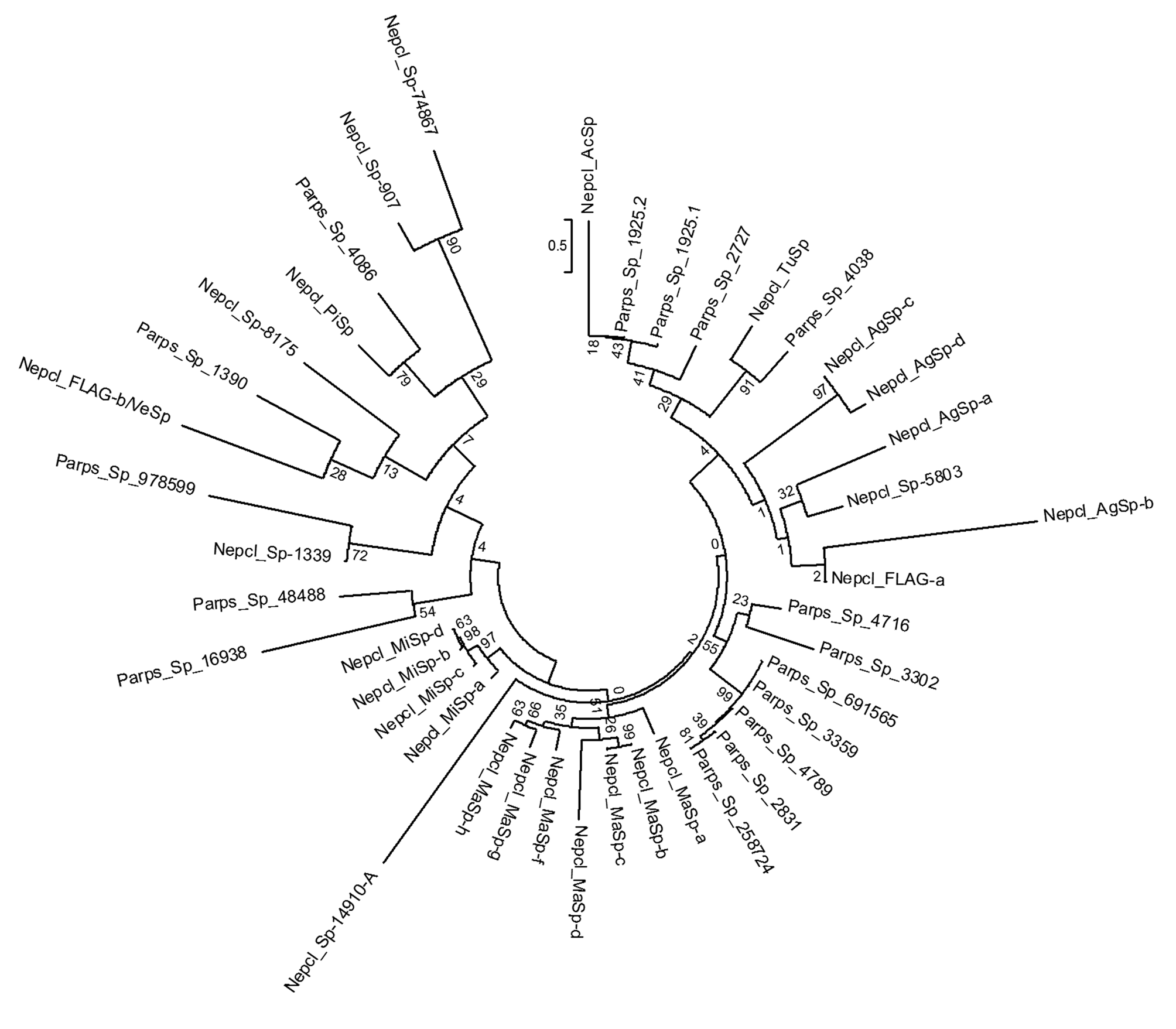

### Figure S5. Neurotoxin U1-lycotoxin-Pp1 nontoxic to mice but toxic to Nillaparvata lugens

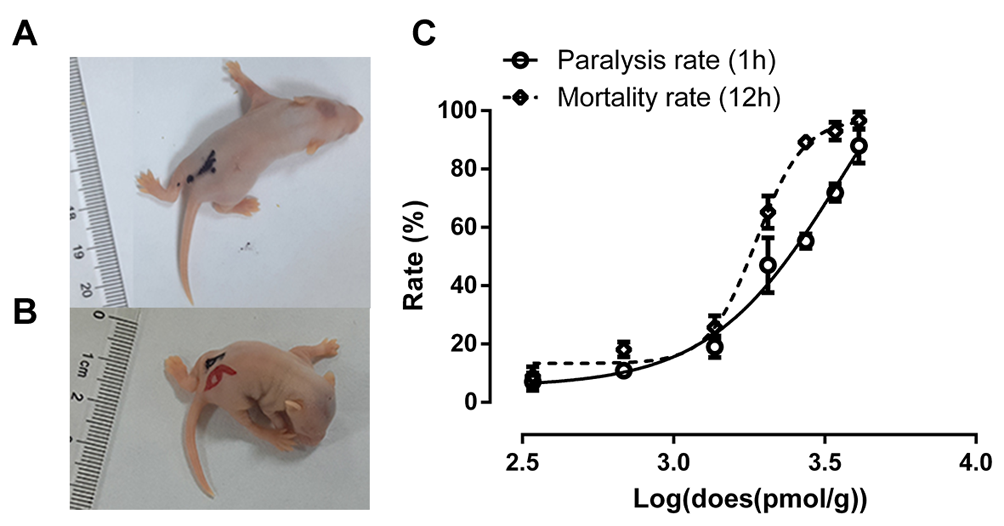

### Figure S6. Linear regression analysis of spider prosoma length and the number of neurotoxins targeting invertebrate prey species.

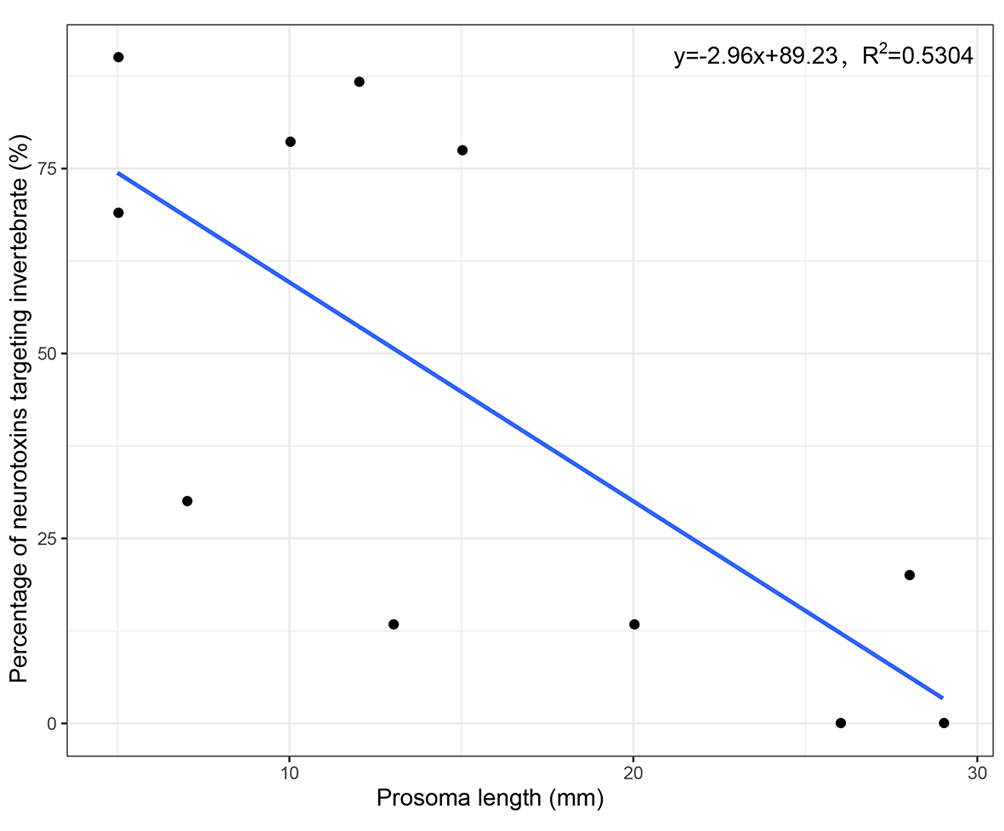
